# Diabetes-associated lung physiology shapes *Klebsiella pneumoniae* infection dynamics

**DOI:** 10.64898/2026.11.12.766331

**Authors:** Krista B. Gunter, Katlyn Todd, James M. Bowen, My T. Le, Kendall K. Wilson, Caitlyn L. Holmes, Karthik Hullahalli, Natasha L. Tilston, Jay Vornhagen

**Affiliations:** Department of Microbiology & Immunology, Indiana University School of Medicine, Indiana University, Indianapolis, USA; Department of Microbiology & Immunology, University of Minnesota, Minneapolis, MN, USA; Department of Microbiology and Immunology, Loyola University Chicago, Maywood, IL USA

## Abstract

Type 2 diabetes mellitus (T2DM) is associated with increased susceptibility to bacterial pneumonia, yet the mechanisms by which diabetic host physiology influences pathogen establishment remain incompletely understood. Here, we investigated how diabetes-associated changes in the lung environment influence *Klebsiella pneumoniae* infection using leptin receptor-deficient (*db/db*) mice. Following acute lung infection, *db/db* mice exhibited increased bacterial burdens compared to nondiabetic controls, despite mounting a robust inflammatory cytokine response. Elevated pulmonary vascular endothelial growth factor concentrations were accompanied by alterations in the bronchoalveolar lavage fluid (BALF) metabolite profile, including enrichment of several amino acids and related metabolites. Consistent with increased nutrient availability, *K. pneumoniae* exhibited enhanced growth in BALF from *db/db* mice and a glutamate-auxotrophic mutant demonstrated improved fitness during pulmonary infection. Using a barcoded *K. pneumoniae* library, we found that *db/db* mice supported significantly larger initial founding populations leading to increased bacterial burden in the lungs. Increased founding populations were also observed in the liver and coincided with proportionally wider infection bottlenecks in the lung. Together, these findings demonstrate that diabetes-associated changes in lung physiology coincide with increased nutrient availability and reduced restrictions on bacterial establishment, highlighting bacterial founding population size as a distinct determinant of pneumonia susceptibility in T2DM.

**Importance:** Type 2 diabetes mellitus increases susceptibility to bacterial pneumonia, but the mechanisms underlying this increased risk remain incompletely understood. Infection severity depends not only on how effectively bacteria replicate but also on how many bacteria successfully establish infection. Here, we demonstrate that the diabetic lung environment is more permissive to *Klebsiella pneumoniae* infection. Diabetes-associated changes in nutrient availability were accompanied by improved fitness of a metabolically constrained bacterial mutant, suggesting that host physiology can alter the metabolic requirements for infection. Using DNA-barcoded *K. pneumoniae*, we further demonstrate that diabetic mice support larger founding populations which increases the downstream bacterial burden in the lungs. These differences were associated with greater bacterial establishment in peripheral tissues, while overall bacterial loads in these tissues remain similar. Collectively, our findings reveal how host metabolic disease influences bacterial fitness and population establishment, providing a framework for understanding increased infection susceptibility in diabetes.

## Introduction

Type 2 diabetes mellitus (T2DM) is a chronic metabolic disease characterized by insulin resistance, impaired insulin secretion, and persistent hyperglycemia^1^. T2DM is associated with increased susceptibility to bacterial infection as well as greater infection-associated morbidity and mortality^2,3^. Although these outcomes are often attributed to impaired immune function, chronic hyperglycemia also causes widespread vascular and metabolic alterations that can affect tissue physiology (reviewed in^4^). The lung is increasingly recognized as a target of diabetic complications, with structural and microvascular abnormalities reported in patients with diabetes^5–8^. However, how these diabetes-associated changes alter the lung environment and influence susceptibility to bacterial infection remains poorly understood.

*Klebsiella pneumoniae* (Kp) is a Gram-negative opportunistic pathogen that causes respiratory, urinary tract, bloodstream, and wound infections^9^. Classical Kp strains are an important cause of hospital-acquired infections, particularly among immunocompromised patient populations, whereas hypervirulent Kp (hvKp) can cause severe community-acquired disease in otherwise healthy individuals and exhibits an increased propensity for invasive and disseminated infection (reviewed in^10^). Kp also represents a significant public health concern because of the increasing prevalence of antimicrobial resistance, with carbapenem-resistant *Enterobacterales,* including Kp, classified as critical-priority bacterial pathogens by the World Health Organization^11^. Because treatment options for Kp are increasingly limited, understanding the host factors that increase susceptibility to severe infection, particularly in at-risk populations such as people with diabetes, is of growing importance.

Diabetes and invasive Kp infection have long been clinically associated. Early reports from Taiwan described Kp liver abscesses with metastatic complications in patients with underlying diabetes, a syndrome subsequently associated with the emergence of hvKp^12,13^. More recent studies have identified diabetes as an important risk factor for Kp infection^14,15^. Despite this well-established epidemiological association, the mechanisms that increase Kp susceptibility and disease severity in the diabetic host remain incompletely understood. While diabetes-associated defects in host immunity may contribute to impaired bacterial clearance, changes to the local tissue environment could also directly influence bacterial growth and fitness.

The pulmonary environment imposes both metabolic and immunological constraints on bacterial infection^16–18^. To establish infection, pathogens must acquire sufficient host-derived nutrients while overcoming barriers that restrict bacterial survival, expansion, and dissemination. Diabetes is associated with pulmonary microvascular dysfunction and increased vascular permeability, changes that may alter the composition and availability of nutrients within the airway^19,20^. These alterations may be particularly relevant for Kp, whose metabolic requirements and fitness vary across host niches. T2DM is also characterized by chronic low-grade inflammation and dysregulated immune responses that can impair effective antimicrobial defense (reviewed in^21^). Whether T2DM alters these metabolic and population-level constraints during Kp infection remains poorly understood.

Here, we tested the hypothesis that T2DM creates a more permissive lung environment that enhances Kp fitness and reduces host-imposed barriers to infection and dissemination. Using a mouse model of T2DM, we examined bacterial burden, lung metabolic and vascular changes, the fitness of a metabolically impaired Kp strain, and bacterial population dynamics during infection.

## Results

### *db/db* mice are more susceptible to Kp lung infection

To measure the impact of T2DM on Kp infection, we compared BKS.Cg-*Dock7^m^* +/+ *Lepr^db/db^*/J mice (“*db/db*” mice^22,23^) to WT C57BL/6J (“*+/+*”) mice. *db/db* mice phenocopy T2DM via development of chronic hyperglycemia, pancreatic β-cell atrophy and hypoinsulinemia. We selected WT C57BL/6J mice as controls as *db/+* heterozygous mice have an intermediate metabolic phenotype^24,25^. For lung infections, we employed our retropharyngeal inoculation method, which replicates acute aspiration pneumonia. We used the hypervirulent Kp strain KPPR1 for these experiments, as this strain is frequently used in animal infection studies and has a wide range of available genetic tools^26^.

Mice were retropharyngeally inoculated with a low (10^4^ colony-forming units [CFUs]) or high (10^6^ CFUs) dose of KPPR1. Higher bacterial burdens were detected in the lungs of *db/db* mice compared to +/+ controls in both dose experiments (**Figure 1A**). We also confirmed that *db/+* mice were less susceptible to Kp pneumonia than *db/db* mice, similar to that of C57Bl/6J (**Figure S1A-B**).

**Figure 1.**
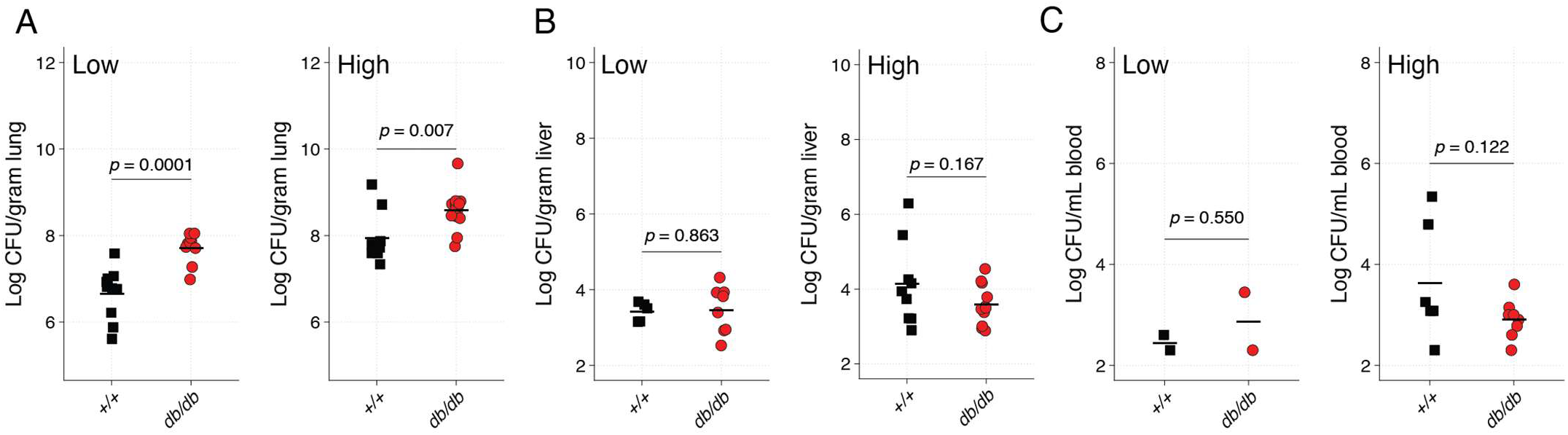
T2DM increases Kp lung burden without a detectable increase in dissemination. Eight-to ten-week-old C57BL/6J (*+/+*, black squares) and BKS.Cg-*Dock7^m^* +/+ *Lepr^db/db^*/J (*db/db*, red circles) mice were retropharyngeally inoculated with either 10^4^ (low dose) or 10^6^ (high dose) CFU of Kp. At 24 hours post-infection, lungs (**A**), livers (**B**), and blood (**C**) were collected, and bacterial burden was quantified. Lung and liver CFU were normalized to tissue weight, and blood CFU were normalized per mL. N = 10 mice per group, each datapoint is an individual mouse, and the black line indicates the group mean. Organs where no CFUs were recovered were excluded from analysis. Statistical comparisons were performed using Student’s *t*-test.

The lung is a common primary site for Kp infection, which can lead to dissemination to the bloodstream, resulting in bacteremia and sepsis (reviewed in^27^). Thus, we also measured bacterial dissemination to the liver and blood. We did not detect differences between liver and blood bacterial loads between *db/db* and *+/+* mice, regardless of dose (**Figure 1B-C**). The two +/+ mice that exhibited high lung burden in the high dose model also exhibited high densities in the liver and blood. Together, these results indicate that the *db/db* mice are more susceptible to acute lung infection than *+/+* mice.

### *db/db* mice exhibit a robust lung cytokine response to Kp infection

Next, we measured cytokine response in mice inoculated with the higher dose of KPPR1 with a Luminex assay using lung homogenate to detect cytokine production relevant to Kp infections. IL-1β (**Figure 2A**), TNFα (**Figure 2B**), and CCL2 (**Figure 2C**) were all significantly elevated in the lungs of *db/db* compared to *+/+* controls. These cytokines were also significantly and positively correlated with lung burden (**Figure 2D-F**). IFNy, IL-6, IL-10, and IL-17A were all elevated due to Kp infection, but no differences were observed between different genotypes (**Figure S2A-D**). IL-6 concentrations were positively correlated with lung bacterial burden, whereas IL-10 and IL-17A concentrations were not. In contrast, IFNy concentrations were negatively correlated with lung bacterial burden (**Figure S2E–H**). These results indicate that *db/db* mice mount a cytokine response to Kp infection that correlates with higher lung CFU. This pattern was also observed in the two *+/+* mice with increased lung bacterial burdens, which had higher cytokine levels.

**Figure 2.**
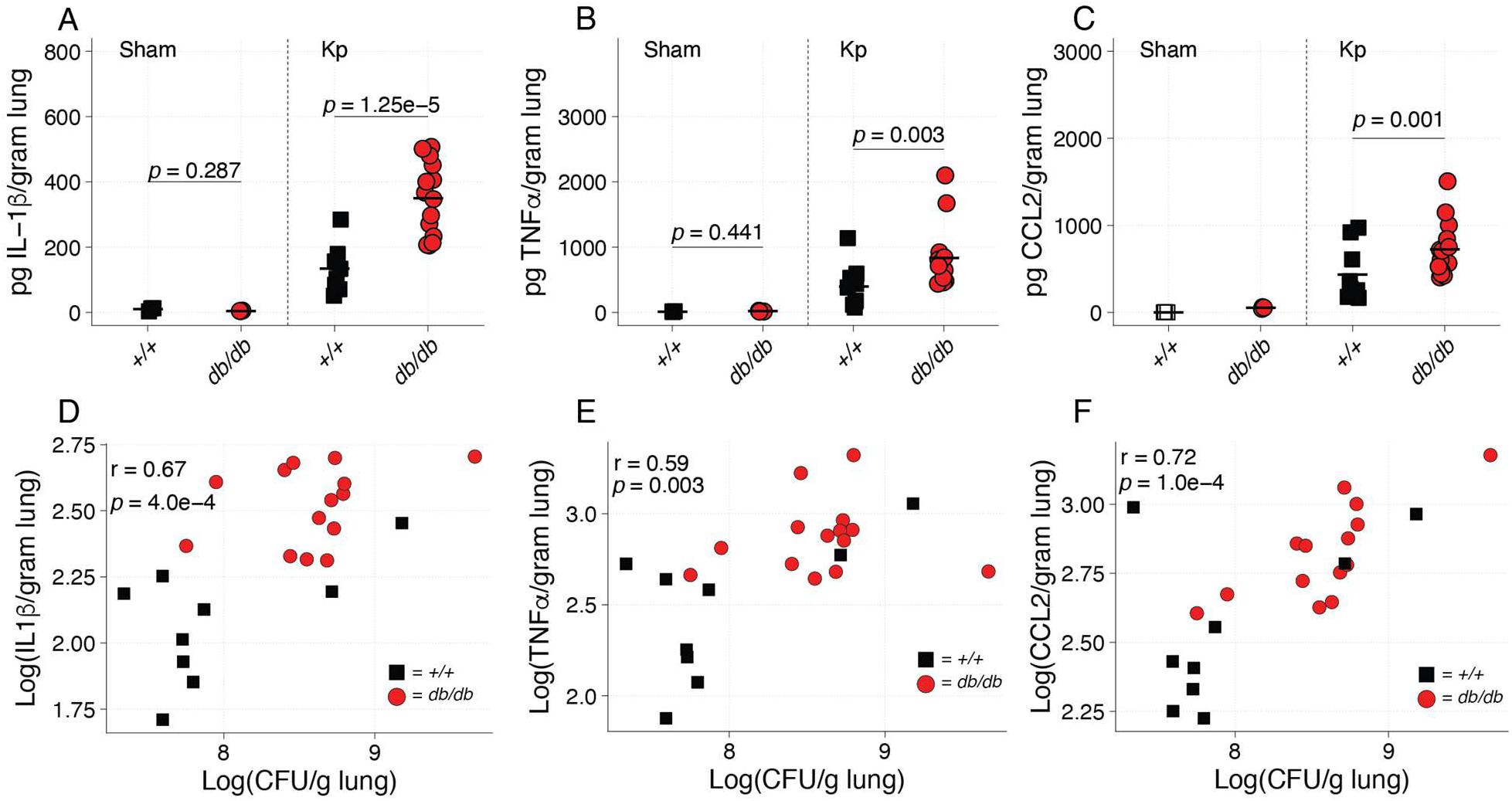
Elevated proinflammatory cytokines in infected *db/db* lungs correlate with bacterial burden. *+/+* (black squares) and *db/db* (red circles) mice were sham inoculated with sterile PBS (N = 3 per genotype) or inoculated with 10^6^ CFU of Kp (N = 10 per genotype). Analyte concentrations were calculated from standard curves and normalized to tissue weight. IL-1β (**A**), TNFα (**B**), and CCL2 (**C**) were higher in infected *db/db* mice than in infected *+/+* mice but did not differ between sham-inoculated groups. Spearman correlation analyses showed associations between lung bacterial burden and IL-1β (**D**), TNFα (**E**), and CCL2 (**F**) concentrations. Points with no fill indicate measurements below the limit of detection. Each datapoint represents an individual mouse and lines indicate group means. For **A-C**, group comparisons were performed using one-way ANOVA followed by Tukey’s post hoc test. Statistical analysis of sham CCL2 levels could not be performed because all +/+ values were below the limit of detection.

### *db/db* mice have elevated levels of lung VEGF

Metabolic diseases, including T2DM, are associated with high vascular permeability due to increases in vascular endothelial growth factor (VEGF^28–30^), which may lead to increases in metabolite availability in the diabetic lung. To test for VEGF changes in our model, we measured VEGF in lung homogenate. VEGF was significantly higher in *db/db* mouse lungs compared to controls at both baseline and during KPPR1 high-dose infection (**Figure 3A**). VEGF concentrations were also positively correlated with lung bacterial burden (**Figure 3B**). Interestingly, VEGF levels increased in infected *db/db* mice, although this difference did not reach statistical significance (Tukey’s HSD *p*= 0.062). No increase was observed in +/+ mice (*p* = 0.661). This pattern suggests that VEGF signaling may be dysregulated in *db/db* mice both at baseline and in response to infection.

**Figure 3.**
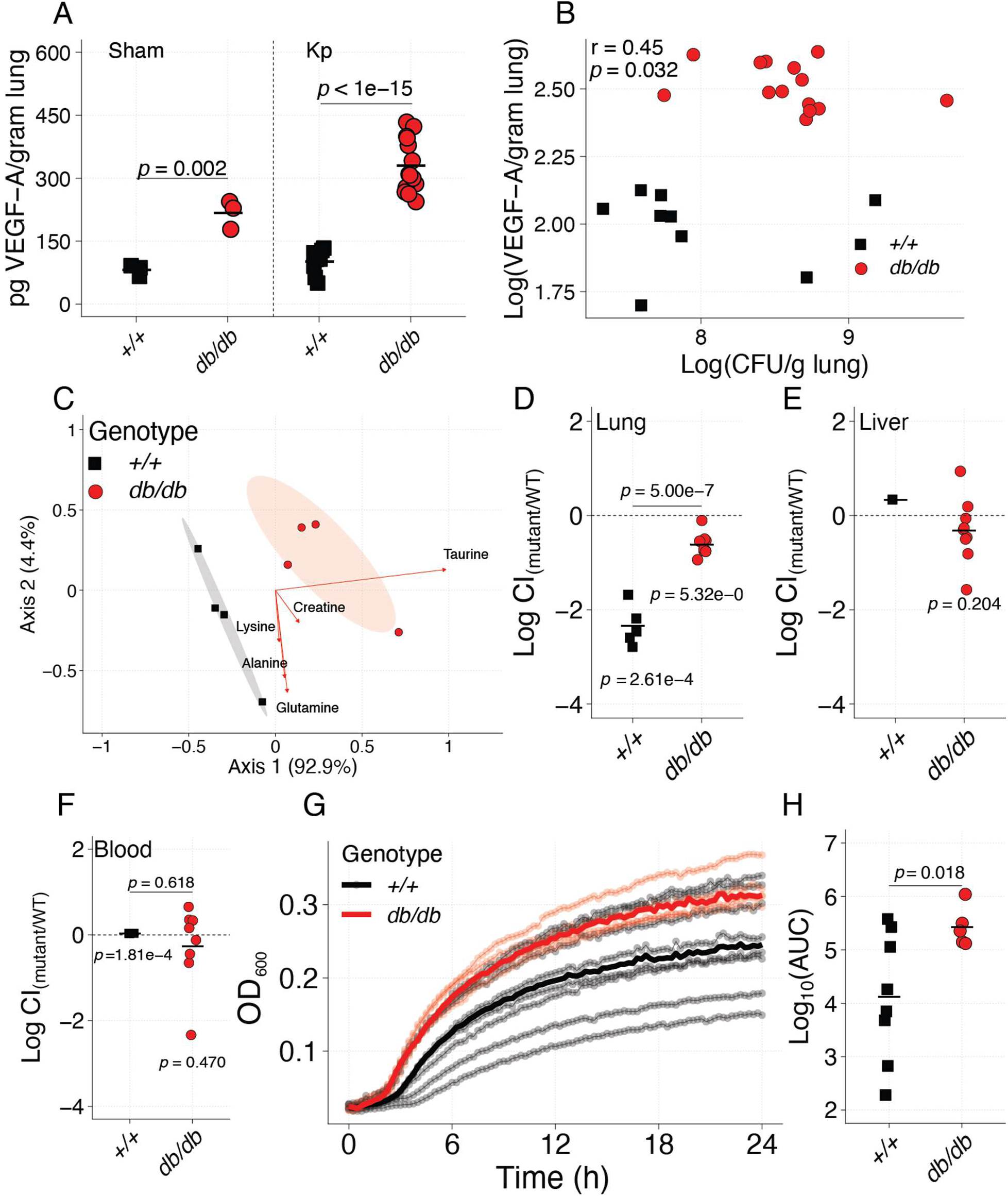
The T2DM lung has an altered amino acid environment that improves the fitness of metabolically constrained Kp. (**A**) VEGF concentrations were measured by Luminex assay in lung homogenates from sham-inoculated (N=3) and Kp-infected (N=15) *+/+* and *db/db* mice. (**B**) Spearman correlation analyses showed associations between lung bacterial burden and concentration of VEGF. (**C**) Principal component analysis of lung amino acid profiles from *+/+* and *db/db* mice (N =4). Amino acid concentrations were normalized to protein content (μg amino acid per μg protein). Each point represents one mouse, and shaded ellipses indicate 70% group ellipses. Red arrows indicate amino acids with an absolute loading greater than 0.2 on either of the first two axes. (**D–F**) *+/+* (black squares) and *db/db* (red circles) mice were retropharyngeally inoculated with a 1:1 mixture of KPPR1S and KPPR1*ΔgltA* mutant. At 24 hours post-infection, lungs (**D**), livers (**E**), and blood (**F**) were collected and plated to quantify each strain and the log_10_ (CFU mutant/CFU wild-type) was calculated. Log_10_ (Competitive index [CI] was compared with a hypothetical mean of zero using a one-sample *t*-test; *+/+* and *db/db* groups were compared using Welch’s *t*-test. (**G**) Growth of KPPR1 in BALF from *+/+* and *db/db* mice normalized by protein content. Transparent points and thin lines represent individual mice, and thick lines show the median OD_600_ at each time point for each genotype. (**H**) Area under the growth curve (AUC) was calculated over 24 hours for each mouse. AUC values were compared between genotypes using Welch’s *t*-test. For **A, D-F, H**, each datapoint represents an individual mouse and lines indicate group means. Datapoints were omitted for organs where no CFUs were recovered.

### The *db/db* lung has altered metabolic environment that facilitates higher Kp growth

Given the positive association between VEGF and vascular permeability, we next hypothesized that the *db/db* lung has increased metabolite availability, which is known to influence Kp infection^31^. Thus, we measured amino acid concentrations in bronchoalveolar lavage fluid (BALF) of uninfected mice. *db/db* mice exhibited a BALF amino acid composition distinct from *+/+* mice (PERMANOVA *p*-value = 0.03, 1,000 permutations, **Figure 3C**). The amino acids most explanatory of composition differences were taurine, creatine, glutamate, glutamine, and lysine, all of which were enriched in the *db/db* compared to *+/+* mice (**Figure 3C**, red arrows). *db/db* mice exhibited modestly higher concentrations of explanatory BALF amino acids compared to *+/+* mice, as well as higher total amino acid concentrations (**Figure S3**).

To confirm the bioavailability of BALF amino acids and their impact on Kp lung infection, we utilized a well-characterized Kp mutant, KPPR1Δ*gltA*. *gltA* encodes citrate synthase, and deletion of this gene renders KPPR1 a glutamate auxotroph^31^. We repeated our high-dose infections using a competition-based approach where KPPR1Δ*gltA* was competed against KPPR1. As expected, KPPR1Δ*gltA* was significantly less fit than its parental strain in the lung of +/+ mice (**Figures 3D, S4A**). Interestingly, the KPPR1Δ*gltA* was significantly more fit in the lungs of *db/db* mice compared to *+/+* mice (**Figures 3D, S4A**), indicating higher bioavailability of glutamate family amino acids in the lungs of *db/db* mice. As expected, GltA was dispensable in the liver and blood of all mouse genotypes (**Figures 3D-F, S4B-C**). These findings indicate that T2DM 1) exacerbates Kp lung infection and 2) increases the bioavailability of amino acids in the lung, potentially through VEGF, thereby opening the lung niche.

Finally, we measured KPPR1 growth *ex vivo* in BALF collected from *+/+* and *db/db* lungs. As expected, KPPR1 growth was more robust in *db/db* BALF compared to *+/+* BALF (**Figure 3G-H**). Interestingly, KPPR1Δ*gltA* was not restored in *db/db* BALF, suggesting that this is not the nutritional compartment that facilitates higher *in vivo*KPPR1Δ*gltA* fitness (**Figure S5**). Collectively, these results indicate that the *db/db* lung has a metabolic environment more favorable for Kp, suggesting a more open lung niche associated with adverse infection outcomes.

### The *db/db* lung sustains a wider bottleneck to infection

Greater CFUs in the lung of *db/db* mice may be a result of a higher founding population or grater replication in the lung niche. To directly measure the breadth of the *db/db* lung niche, we employed Sequence-Tagged Analysis of Microbial Populations with resampling (STAMPR) using a fitness-neutral barcoded KPPR1 library^32^. To this end, we repeated our high-dose acute lung infection studies. Consistent with previous results, *db/db* mice had increased lung burden compared to *+/+* mice (**Figure 4A**). Interestingly, the number of unique founders (N_s_) was increased in the *db/db* lung compared to the *+/+* lung (**Figure 4B-C**), indicating that diabetic lungs are more vulnerable to initial seeding by Kp. Normalization of N_s_ to CFUs provides a measure of net replication, which did not differ between the genotypes (**Figure 4D**), indicating diabetic lung tissue does not enhance the ability for Kp clones to replicate. Collectively, these results indicate that the *db/db* mice primarily differ in the number of bacteria that successfully contribute to the acute pneumonia, rather than producing a readily detectable increase in expansion per founder.

**Figure 4.**
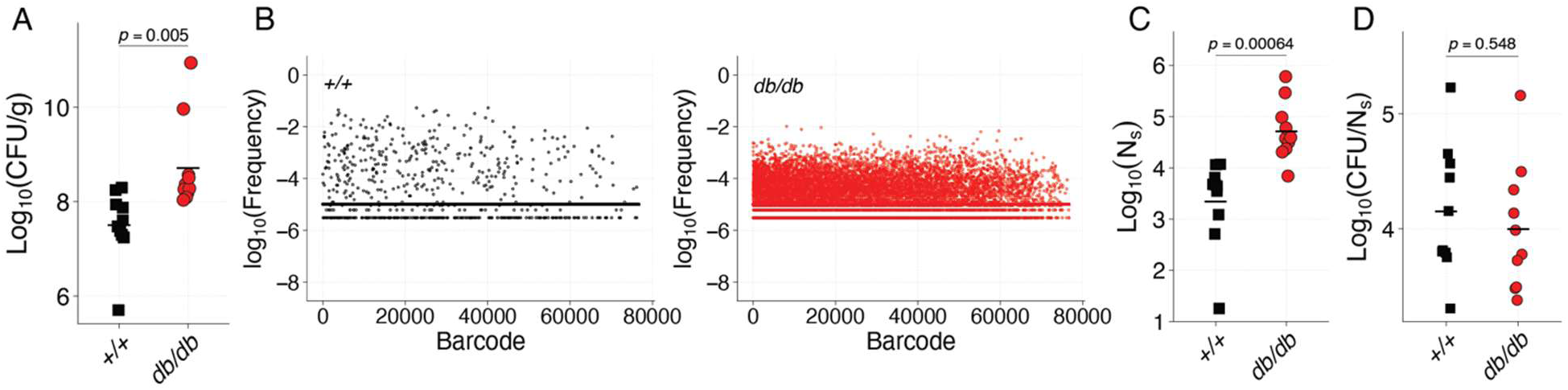
A larger bacterial founding population establishes infection in T2DM lungs. *+/+* (black squares) and *db/db (*red circles) mice were retropharyngeally inoculated with a library of approximately 40,000 barcoded Kp strains. Lungs were collected at 24 hours post-infection to quantify bacterial burden and analyze barcode populations using STAMPR. (**A**) Lung bacterial burden, (**B**) representative plots for barcode frequency in lung tissue for +/+ (black) and *db/db* (red) from a single mouse per genotype (**C**) estimated founding population size (N_s_), and (**D**) CFU/N_s_. For **A, C,** and **D** each point represents an individual mouse, N=10. For **B**, each point represents individual barcoded Kp mutant. For **C-D**, Welch’s *t*-test used for statistical analysis.

### Disseminated Kp populations reflect the lung founding population in *db/db* mice

Unlike our original infection studies, our STAMPR studies showed modest, but non-significant, increase in the bacterial burden in the liver of *db/db* mice (**Figure 5A**). The number of unique founders (N_s_) was significantly greater in the *db/db* liver, indicating that more bacterial lineages contributed to the hepatic population (**Figure 5B**). Like the lung, normalization of N_s_ to CFUs did not differ between genotypes, suggesting that the increased number of founders was not accompanied by a detectable difference in clonal expansion (**Figure 5C**). Additionally, these values were very close to 0, suggesting that Kp does not rapidly replicate in the liver in our model. Thus, in *db/db* mice, a greater number of clones arrive in the lungs and then can escape in the liver, however the liver environment does not favor grater replication of these clones.

**Figure 5.**
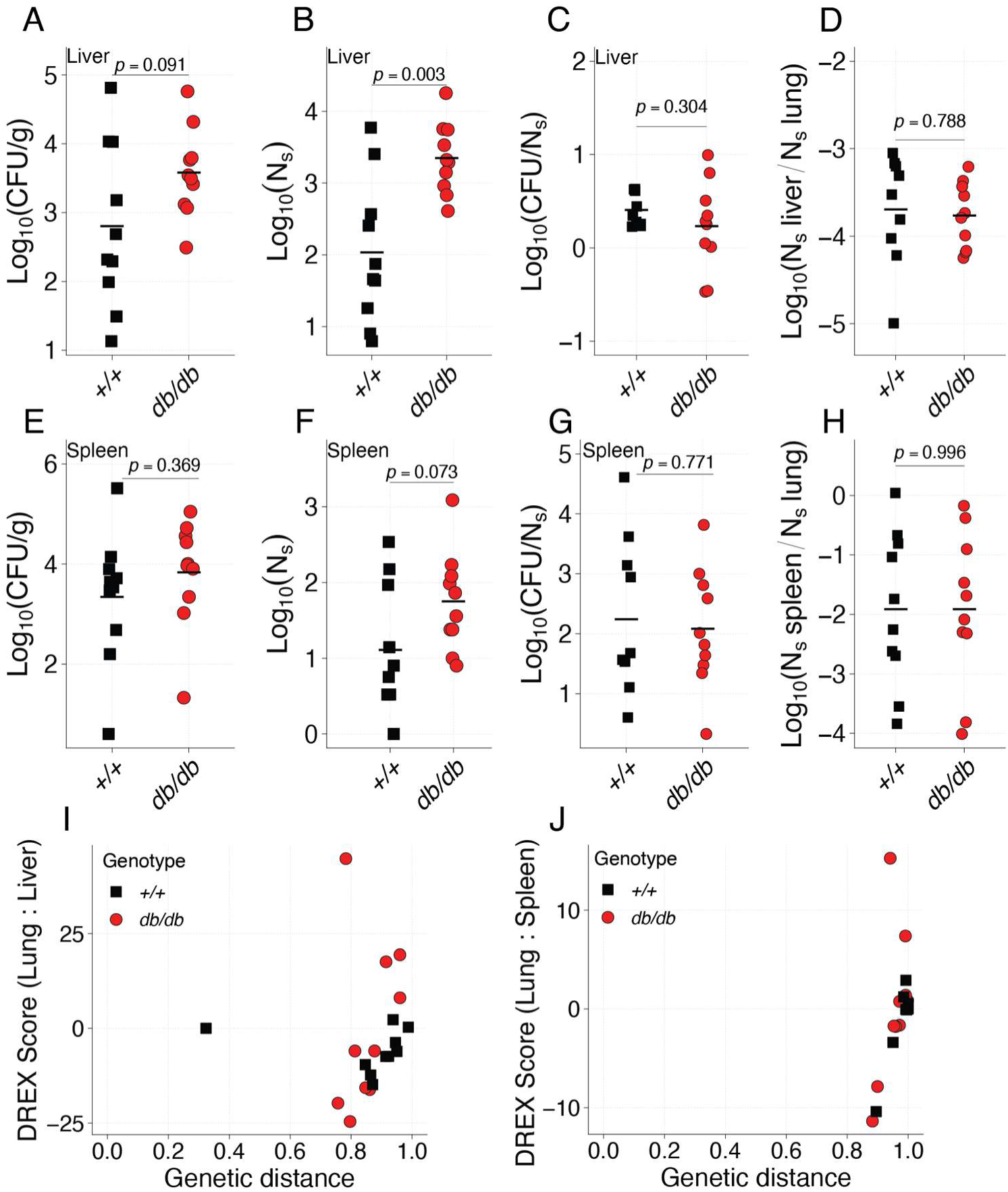
T2DM increases bacterial founding population size in the liver and spleen without increasing bacterial burden. *+/+* (black squares) and *db/db* (red circles) mice were retropharyngeally inoculated with a library of approximately 40,000 barcoded Kp strains. Livers (**A–D**) and spleens (**E–H**) were collected at 24 hours post-infection to quantify bacterial burden and analyze barcode populations using STAMPR. Bacterial burden in the liver (**A**) or spleen (**E**). Founding population size (N_s_) in the liver (**B**) and spleen (**F**). Bacterial burden relative to founding population size, calculated as log_10_ (CFU/N_s_), in the liver (**C**) or spleen (**G**). Genotypes were compared using Welch’s t test. Genetic distance (GD) and distance from randomly sampled expectations (DREX) scores were calculated between lung and liver (**D**) and lung and spleen (**H**) bacterial populations from individual mice. Each point represents one mouse (N=10 per group).

To determine whether this reflected a wider dissemination bottleneck, we normalized hepatic N_s_ to the corresponding lung N_s_ within each animal to determine whether high liver founders were those with high lung founders (**Figure 5D**). This ratio did not differ between genotypes, suggesting that diabetes does not disproportionally increase the number of clones exiting the lung. Rather, the increase in diversity in the lung leads to a proportional increase in the number of clones that exit the lung and reach the liver. Collectively, these findings indicate that the *db/db* liver supports larger founding populations following lung infection. This is consistent with increased systemic establishment associated with the expanded pulmonary founding population.

We similarly examined dissemination to the spleen, where *db/db* mice exhibited a modest but nonsignificant increase in bacterial burden compared to *+/+* mice (**Figure 5E**). Splenic Ns were also increased in *db/db* mice, although this difference did not reach statistical significance (**Figure 5F**). As observed in the lung and liver, normalization of Ns to CFUs did not differ between genotypes, suggesting comparable clonal expansion relative to the number of bacterial founders (**Figure 5G**). Additionally, these values were greater than 0, suggesting that, unlike the liver, Kp replicated more rapidly in the spleen. Interestingly, normalization of splenic N_s_ to lung N_s_ revealed no increase in relative systemic founding population size in *db/db* mice, suggesting that the increased lung founding population did not result in a proportionally greater establishment of splenic founders (**Figure 5H**). Together, these findings indicate that, although diabetes is associated with increased bacterial establishment in the lung and liver, the spleen retains a comparatively restrictive bottleneck that limits the number of bacterial lineages contributing to systemic infection.

Finally, to determine whether increased systemic founding populations in *db/db* mice were associated with altered patterns of bacterial dissemination, we compared barcode distributions between organs. Genetic distances (GD), where values approaching 1 indicate highly dissimilar populations, were generally high between the lung and both peripheral organs, as well as between the liver and spleen, with no consistent genotype-dependent differences (**Figure S6**). Because increased founding population sizes can influence barcode similarity independently of dissemination, we additionally calculated the distance from randomly sampled expectations (DREX). DREX scores were used to assess whether barcode similarity between the lung and other organs differed from what would be expected under random sampling. Negative scores indicate greater-than-expected similarity, whereas positive scores indicate lower-than-expected similarity. The lung–liver DREX and lung–spleen DREX (Welch’s *t*-test, *p* = 0.171, *p* = 0.123) in *+/+* mice compared to *db/db* mice did not significantly differ between genotypes (**Figure 5I-J**). Collectively, these results suggest that the increased founding populations in *db/db* mice are not accompanied by greater sharing of expanded pulmonary clones but instead support distinct bacterial population structures across infection sites.

## Discussion

In this study, we found that T2DM increased Kp lung burden and altered the pulmonary environment in ways that favored bacterial growth and establishment. T2DM mice had higher baseline levels of VEGF, a regulator of vascular permeability. To investigate whether the lung nutrient environment was also altered, we measured lung amino acids and found that T2DM lungs had higher total concentrations and a distinct amino acid profile compared with wild-type lungs. Consistent with a more favorable nutrient environment, the metabolically constrained *gltA* mutant grew better in T2DM lungs than in wild-type lungs. Using STAMPR, we also found that a larger bacterial founding population established infection in T2DM lungs. Although susceptibility to infection in T2DM is often attributed to altered immune defenses, our findings suggest that host metabolic disease also may create greater opportunities for bacterial establishment and growth.

Our data contribute to a growing understanding that the lung nutritional environment is an important determinant of pathogen fitness and infection severity. Previously, we demonstrated that Kp lung fitness depends on the ability to access and utilize metabolites in the local environment, a property we described as metabolic flexibility^31^. Here, we found that the fitness advantage conferred by metabolic flexibility was reduced in the diabetic lung, as demonstrated by the increased fitness of the metabolically constrained *gltA* mutant. Given the extensive metabolic heterogeneity among Kp strains^33^, these findings raise the possibility that individuals with diabetes may be susceptible to a broader range of Kp strains with distinct metabolic capabilities. We hypothesize that this increased metabolic permissiveness is driven, at least in part, by VEGF-associated changes in lung vascular permeability that increase the availability of circulating nutrients. Alterations in VEGF signaling has been previously shown in diabetic patients to affect disease pathogenesis (reviewed in^34^). Consistent with this model, vascular leakage has previously been shown to increase lung nutrient availability and promote secondary bacterial pneumonia following influenza infection^35–38^, while diabetes-associated increases in airway glucose can enhance bacterial growth during respiratory infection^39^. Moreover, our previous work demonstrated comparable WT and *gltA* fitness in serum, suggesting that access to circulating nutrients could partially alleviate the metabolic constraints imposed on Kp during lung infection^31^. Interestingly, although BALF from *db/db* mice supported increased WT Kp growth, it did not restore *gltA* fitness, suggesting that the sampled airway compartment does not fully recapitulate the metabolic environment responsible for the *in vivo* phenotype. This discrepancy may reflect spatial heterogeneity in nutrient availability between pulmonary compartments or infection-associated changes not captured by BALF collected from uninfected mice. For example, increased taurine in the BALF may drive increased Kp growth, whereas glutamate-family amino acids that drive *gltA* may be more available at sites of increased vascularization. Future studies should determine whether VEGF-dependent changes in vascular permeability contribute causally to increased nutrient availability and identify the specific lung microenvironments that support Kp fitness.

The STAMPR approach used in this study independently supports the conclusion that the diabetic lung is more permissive to Kp infection. Our data demonstrate that increased pulmonary bacterial burden in *db/db* mice is associated with a larger founding population without a detectable increase in average clonal expansion, suggesting that more bacteria successfully contribute to the established infection^32^. Several nonexclusive mechanisms could explain this difference. First, increased nutrient availability may promote bacterial survival during early infection, relaxing the effective founding bottleneck by allowing more bacteria to establish themselves in the pulmonary environment. Second, diabetes-associated defects in innate immunity may reduce the efficiency of early bacterial clearance. Indeed, previous studies have demonstrated impaired granulocyte recruitment and altered inflammatory signaling during Kp pneumonia in diabetic mice^40^. Although our data demonstrate a robust cytokine response in *db/db* mice, we did not assess immune-cell recruitment or the effectiveness of antimicrobial responses. Third, diabetes-associated alterations in pulmonary tissue architecture and epithelial or vascular barrier function may modify bacterial access to, and survival within, local tissue compartments^8,41^. Importantly, these processes are unlikely to act independently, as nutrient availability, immune restriction, and tissue architecture collectively shape the conditions under which bacteria establish infection. Together, these findings raise the possibility that diabetes-associated changes in the lung environment alter not only the metabolic requirements for bacterial fitness but also the probability that individual bacteria successfully establish infection.

Changes in lung susceptibility may also have important consequences for disseminated infection. We observed a larger hepatic founding population in *db/db* mice, although the ratio of hepatic to pulmonary N_s_ did not differ between genotypes. This suggests that increased hepatic establishment largely reflects the expanded pulmonary founding population, rather than a genotype-specific relaxation of the lung-to-liver dissemination bottleneck. Indeed, tissue-resident immune defenses, including hepatic Kupffer cells, have been shown to restrict circulating Kp, demonstrating that bacterial survival at secondary sites depends on local host defenses^42^. Interestingly, our GD and DREX analyses did not demonstrate increased sharing of dominant lung clones in *db/db* mice, despite their larger founding populations. These findings are broadly consistent with the previously described pattern of direct Kp dissemination, wherein bacteria reach secondary sites without extensive expansion at the primary infection site, rather than metastatic dissemination driven by the spread of dominant primary site clones^32^. However, early dissemination followed by independent population expansion or clearance could produce similar patterns, and the relative contributions of these processes remain unresolved. The clinical implications of these findings are particularly relevant given that lung infection is associated with increased mortality among patients with diabetes and Kp bacteremia^15^. Together, our findings suggest that diabetes-associated increases in pulmonary establishment may contribute to systemic infection without necessarily altering the relative barriers to dissemination, emphasizing the importance of primary-site susceptibility in determining infection outcomes.

While this study advances our understanding of Kp lung infection during diabetes, several limitations remain. First, although elevated VEGF, altered lung nutrient availability, and increased bacterial fitness support a model of enhanced metabolic permissiveness, the causal relationships between VEGF signaling, vascular permeability, nutrient availability, and bacterial establishment remain to be established. Moreover, altered immune function may contribute to the increased founding population in *db/db* mice, and the relative contributions of metabolic and immunological changes remain unresolved. Second, our studies employed leptin receptor-deficient *db/db* mice, and leptin signaling itself influences lung antimicrobial defenses, potentially contributing to susceptibility independently of diabetes-associated metabolic dysfunction^43^. Finally, our infection studies were limited to a single acute timepoint and a single Kp strain, restricting our ability to resolve the temporal dynamics of bacterial establishment and determine the generalizability of these findings across the metabolic diversity of Kp. Future studies should aim to distinguish the relative contributions of these mechanisms and determine whether correcting diabetes-associated metabolic or vascular abnormalities can restore pulmonary restriction of bacterial infection. Together, our findings demonstrate that diabetes-associated changes in lung physiology influence both bacterial metabolic fitness and the number of bacteria that successfully establish infection, identifying host metabolic state as an important determinant of the physiological barriers restricting bacterial pathogenesis.

## Materials and methods

### Ethics statement

This study was performed strictly in accordance with the recommendations in the Guide for the Care and Use of Laboratory Animals^44^. The Indiana University School of Medicine Institutional Animal Care and Use Committee approved this research (Protocols 22114 and 25101; PI: Vornhagen).

### Materials, media, and bacterial strains

The Kp strain KPPR1 was used for lung infection experiments. Additional strains in this study include streptomycin-resistant KPPR1 (KPPR1S), the isogenic KPPR1Δ*gltA* mutant^31^, and the KPPR1 STAMPR library^32^. All strains were cultured in Luria-Bertani (Miller formulation) broth at 37°C with shaking or on Luria-Bertani agar at 27°C with 40 μg/ml rifampicin (KPPR1, STAMPR), 40 μg/ml kanamycin (KPPR1Δ*gltA*), or streptomycin 500 µg/mL (KPPR1S). All materials and reagents were purchased from Millipore-Sigma (St. Louis, MO) or Thermo Fisher Scientific (Waltham, MA).

### Murine model of infection

Eight-to 10-week-old mice (The Jackson Lab strains 000642 and 000664) were used for all murine models, as this time point corresponds to the spontaneous onset of T2DM-like disease^22^. Lung infections were performed as previously described with no modifications^31^. Each Kp strain was cultured overnight with appropriate antibiotics. Bacteria were then pelleted, resuspended, and diluted in sterile phosphate-buffered saline (PBS). For competitive studies, KPPR1 and the isogenic KPPR1Δ*gltA* mutant were mixed 1:1. Mice were anesthetized with isoflurane, inoculated in the pharynx with a total inoculum of 10^4^ or 10^6^ CFU in 50 mL or an equal volume of PBS (Sham), and monitored until ambulatory. 24 hours post-inoculation, mice were euthanized by CO_2_ asphyxiation, and tissues were collected, weighed, and homogenized in sterile PBS. Tissue homogenates were dilution plated on selective media to determine bacterial load and competitive index. For STAMPR studies, the remaining organ homogenate was plated onto LB agar with kanamycin to recover the total number of barcodes within each site, and plates were incubated overnight at 27 °C. The following day, bacterial lawns were scraped and mixed with sufficient water to create a turbid suspension, keeping each mouse and tissue separate. 100 µL was removed from each sample and boiled to extract bacterial DNA. The remaining bacteria collected from lawn scraping were stored in 20% glycerol at −80 °C. Mice were monitored for signs of distress and euthanized at predetermined time points. No blinding was performed between experimental groups.

### Lung analyte measurements

For cytokine analysis, lung homogenates were subjected to Luminex analysis. A custom Luminex assay including IL-1β, IL-6, IL-10 and IL-17A, CCL2, IFNγ, TNFα, and VEGF-A (assay ID: MX2XAXN) was assembled and performed using a Luminex 200 instrument (Thermo Fisher Scientific) per manufacturer’s instructions. 50 events were collected for each sample and standard.

For metabolite measurement, bronchoalveolar lavage fluid (BALF) was collected as previously described^31^. Uninfected mice were euthanized by CO_2_ asphyxiation. Following euthanasia, a small incision was made in the trachea, and polyethylene tubing (external diameter 0.965 mm, internal diameter 0.58 mm, BD, Franklin Lakes, NJ) with a 23-gauge luer-stub adaptor was inserted and secured with 4–0 silk suture (Ethicon, Somerville, NJ). 2 mL sterile PBS was used to flush the lungs. BALF was centrifuged at 21,130 x *g* for 15 min at 4°C to pellet contaminating bacteria and supernatants were stored at -80 °C for amino acid analysis. BALF amino acid concentrations were quantified using liquid chromatography–tandem mass spectrometry as previously described^45^. BALF amino acid concentrations were normalized to total protein concentration, which was measured by BCA Protein Assay (Thermo Fisher Scientific) per manufacturer’s instructions. Only amino acids detected in >50% of BALF samples were used for final analysis. Differences in metabolome structure were assessed by PERMANOVA (1,000 permutations) using Euclidean distance as the dissimilarity metric from the vegan package, v. 2.6-2.

To assess the capacity of BALF to support bacterial growth, BALF was collected from uninfected mice as described above. For growth curves, 1 µL of an overnight KPPR1 culture was inoculated into normalized BALF. Cultures were incubated at 37 °C in a BioTek Synergy plate reader, and optical density at 600 nm (OD_600_) was measured every 15 minutes for 24 hours. For competition assays, normalized BALF was inoculated with a 1:1 mixture of KPPR1S and KPPR1 Δ*gltA* and incubated at 37 °C for 24 hours. Cultures were diluted and spot-plated on streptomycin (500 µg/mL) and kanamycin (40 µg/mL) plates to quantify KPPR1S and KPPR1 Δ*gltA*, respectively. The competitive index (CI) was calculated as output KPPR1Δ*gltA* CFUs/mL divided by KPPR1S CFUs/mL divided by input KPPR1Δ*gltA* CFUs/mL divided by KPPR1S CFUs/mL.

### DNA sequencing and STAMPR analysis

DNA sequencing and STAMPR analysis were performed as previously described^32^. Briefly, bacterial barcodes were amplified by PCR and sequenced on an Illumina platform, and sequencing reads were mapped to corresponding reference barcode library and analyzed using the STAMPR pipeline to estimate founding population size, genetic distance, and fractional relatedness.

### Statistical Analysis

All animal studies were replicated at least twice with biological sex evenly distributed. Competitive indices were log_10_ transformed, and a one-sample *t-*test was used to determine significant differences from a hypothetical value of 0. All colony-forming unit values were log_10_ transformed for analysis. A *p*-value of less than 0.05 was considered statistically significant for the above experiments. Welch’s *t*-test or ANOVA followed by Tukey’s multiple comparisons post-hoc test was used to determine significant differences between groups. Analysis was performed using RStudio 2021.09.0+351 “Ghost Orchid” release for macOS with R v. 4.4.0.

### Data availability

All raw data for this study are in **Dataset S1,** and analysis code are available at 10.5281/zenodo.24173614. The raw input data for DREX calculation are available at 10.5281/zenodo.17573028.

## Acknowledgments

The authors would like to acknowledge the members of the Tilston and Vornhagen labs for their feedback on this study.

## Contributions

Conceptualization: CLH, JV

Methodology: KH, CLH, JV

Investigation: All authors

Visualization: KG

Funding acquisition: CLH, JV

Project administration: JV

Supervision: JV

Writing – original draft: KG, KT, JV

Writing – review & editing: All authors

## Use of AI

ChatGPT was used after drafting this manuscript for grammar and clarity improvements.

## Supplementary data

**Supplemental Figure 1.**
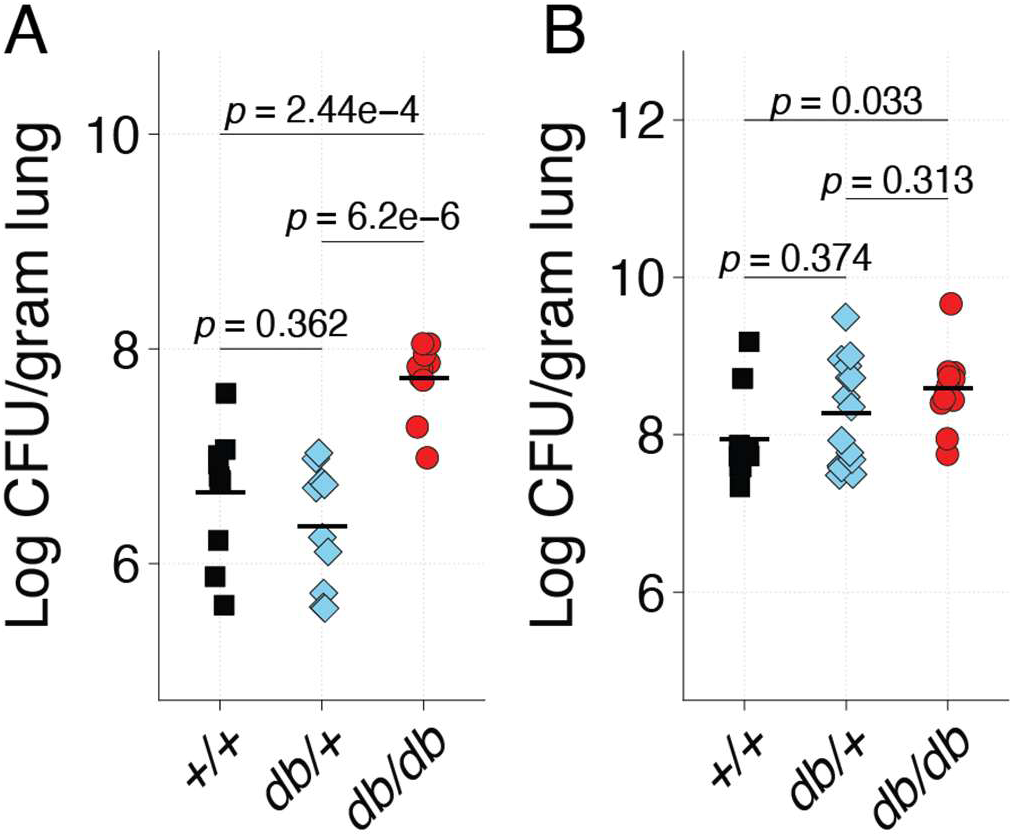
Kp lung burden does not differ in heterozygous or C57BL/6J mice. Six-to eight-week-old C57BL/6J (*+/+*, black squares) and heterozygous BKS.Cg-*Dock7^m^* +/+ *Lepr^db/db^*/J (*db/+*, purple diamonds) mice were retropharyngeally inoculated with either 10^4^ (low dose) (**A**) or 10^6^ (high dose) (**B**) CFU of Kp. At 24 hours post-infection, lungs were collected, and bacterial burden was quantified. Each point represents an individual mouse (N=10 per group), and black lines indicate the group mean. Statistical comparisons were performed using one-way ANOVA followed by Tukey’s HSD post hoc test.

**Supplemental Figure 2.**
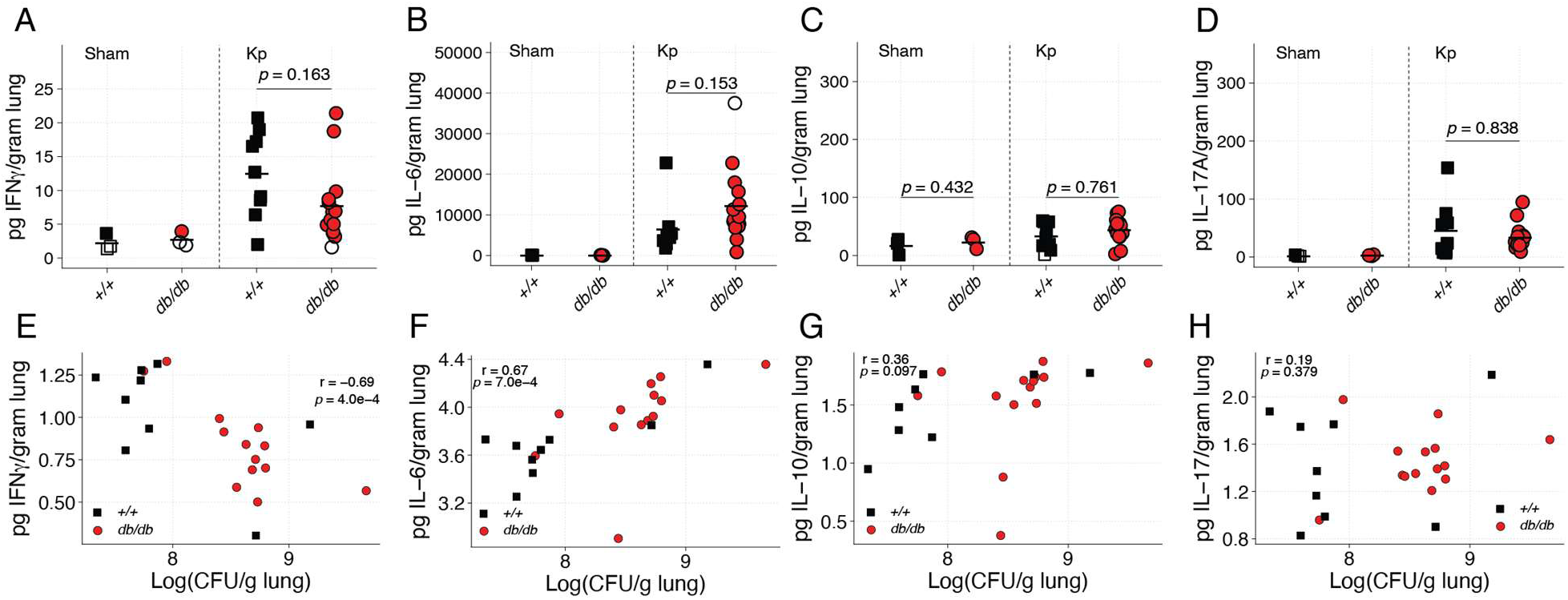
Cytokines that respond to Kp lung infection independent of genotype. *+/+* (black squares) and *db/db* (red circles) mice were sham inoculated with sterile PBS (N = 3 per genotype) or inoculated with 10^6^ CFU of Kp (N = 10 per genotype). Analyte concentrations were calculated from standard curves and normalized to tissue weight. IFNy (**A**), IL-6 (**B**), IL-10 (**C**), and IL-17A (**D**). Spearman correlation analyses showed associations between lung bacterial burden and IFNy (**E**), IL-6 (**F**), IL-10 (**G**), and IL-17 (**H**) concentrations. Points outlined in light blue indicate measurements the limit of detection. Each datapoint represents an individual mouse and lines indicate group means. For **A-D**, group comparisons were performed using one-way ANOVA followed by Tukey’s post hoc test.

**Supplemental Figure 3.**
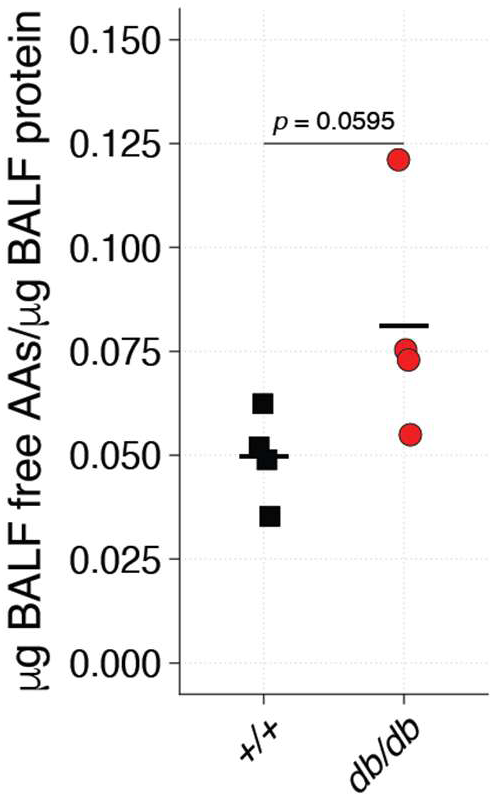
Concentration of free amino acid in BALF. BALF was collected from *+/+* (black squares) and *db/db* (red circles) from uninfected mice to measure free amino acids using liquid chromatography-tandem mass spectrometry (N = 4 per genotype). Concentration of free amino acids was normalized to total protein content. Each point represents an individual mouse, and black lines indicate the group mean. Welch’s *t*-test was used for statistical analysis.

**Supplemental Figure 4.**
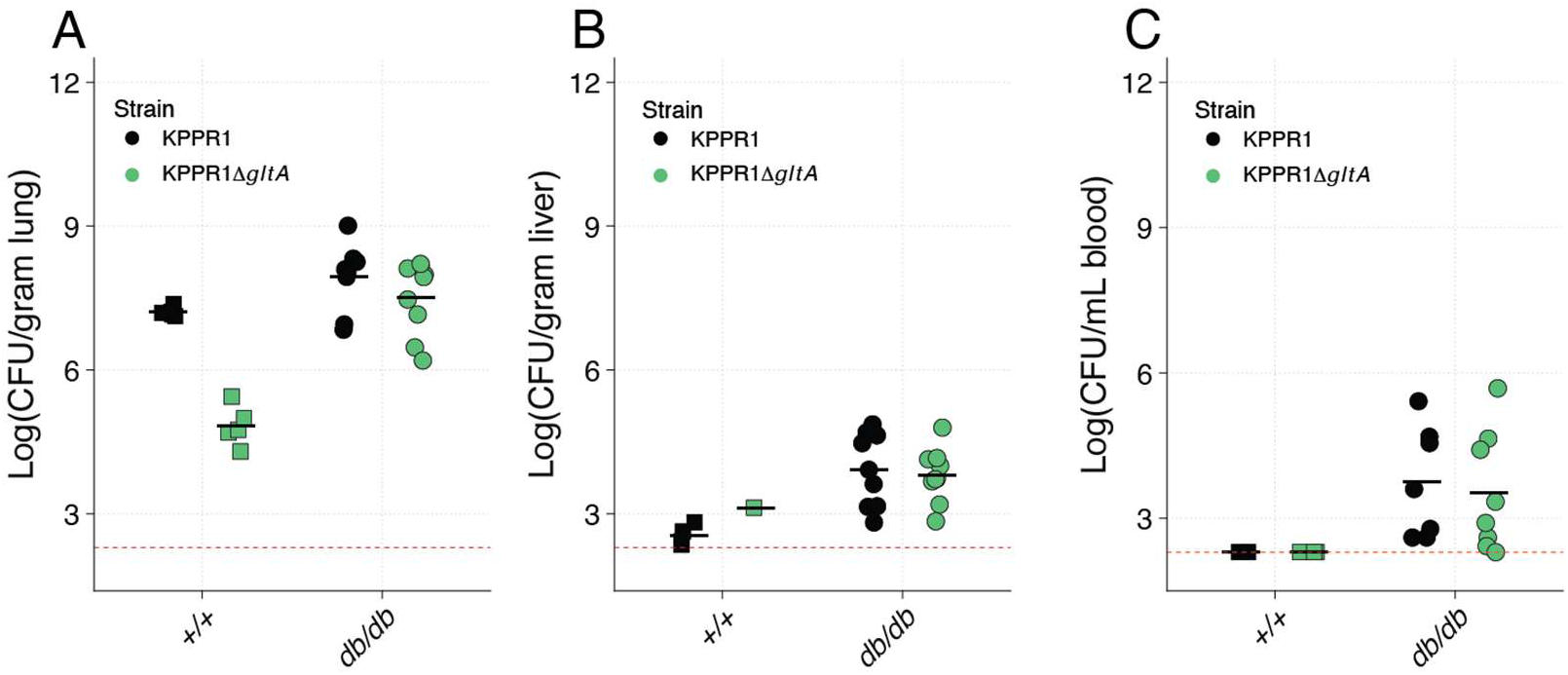
CFU quantification of *gltA* mutant. *+/+* and *db/db* mice were retropharyngeally inoculated 1:1 KPPR1 (black) and the KPPR1Δ*gltA* mutant (green) to a total concentration of 10^6^ CFU/mouse. 24 hours post-inoculation lung (**A**), liver (**B**), and blood (**C**) CFUs were quantified on selective media to determine bacterial load per strain. Each point represents an individual mouse, and black lines indicate the group mean. Dashed red line indicates the limit of detection for assay.

**Supplemental Figure 5.**
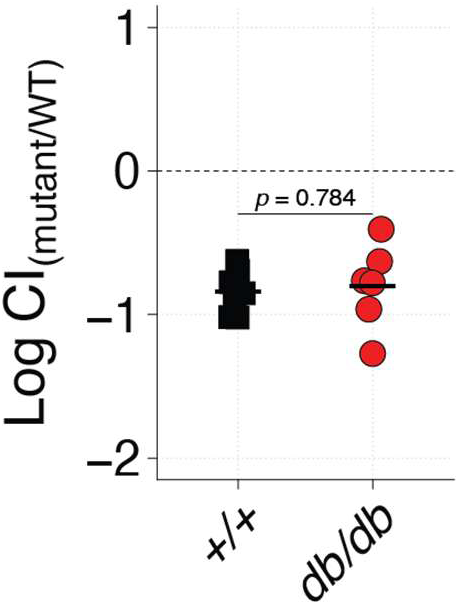
d*b/db* BALF does not restore *gltA* fitness. BALF from *+/+* (black squares) and *db/db* (red circles) mice normalized by protein content was inoculated 1:1 with KPPR1 and KPPR1Δ*gltA*. 24 hours post infection CFUs were quantified on selective media to determine CFU per strain and calculate competitive index. Each point represents an individual mouse, and black lines indicate the group mean. Statistical analysis was performed using Welch’s *t*-test.

**Supplemental 6.**
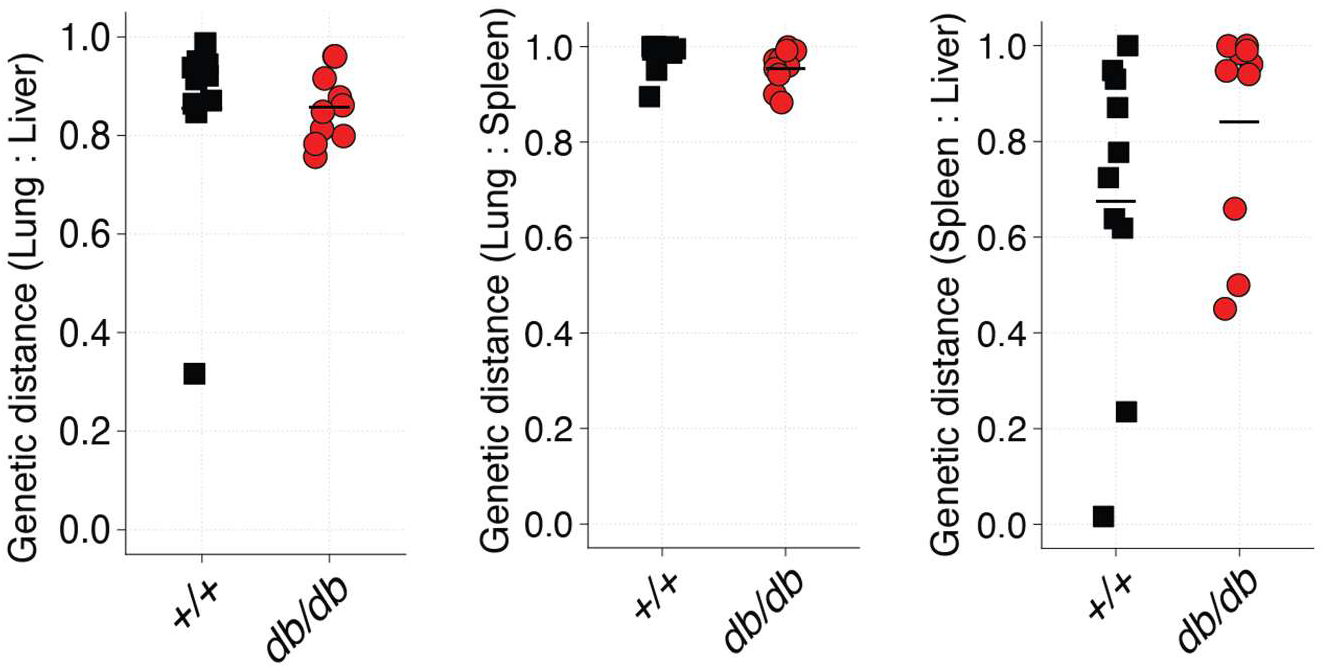
Mouse genotype is not associated with genetic distance. Genetic distances between lung and liver, lung and spleen, and liver and lung.

